# Relationship Between Irregularities of the Nuclear Envelope and Mitochondria in HeLa cells Observed with Electron Microscopy

**DOI:** 10.1101/2023.11.14.567016

**Authors:** D. Brito-Pacheco, C. Karabağ, C. Brito-Loeza, P. Giannopoulos, C.C. Reyes-Aldasoro

**Affiliations:** CLIR, Facultad de Matemáticas, Universidad Autónoma de Yucatán, Mexico; School of Science and Technology, City, University of London, UK; School of Computing and Digital Media, London Metropolitan University, UK

**Keywords:** Segmentation, HeLa Cells, Mitochondria, Invaginations

## Abstract

This paper describes a methodology to analyse the complexity of HeLa cells as observed with electron microscopy, in particular the relationship between mitochondria and the roughness of the nuclear envelope as reflected by the invaginations of the surface. For this purpose, several segmentation mitochondria algorithms were quantitatively compared, namely: Topology, Image Processing, Topology and Image Processing, and Deep Learning, which provided the highest accuracy. The invaginations were successfully segmented with one image processing algorithm. Metrics were extracted for both structures and correlations between the mitochondria and invaginations were explored for 25 segmented cells. It was found that there was a positive correlation between the volume of invaginations and the volume of mitochondria, and negative correlations between the number and the mean volume of mitochondria, and between the volume of the cytoplasm and the aspect ratio of mitochondria. These results suggest that there is a relationship between the shape of a cell, its nucleus and its mitochondria; as well as a relationship between the number of mitochondria and their shapes. Whilst these results were obtained from a single cell line and a relatively small number of cells, they encourage further study as the methodology proposed can be easily applied to other cells and settings.

Code and data are freely available. HeLa images are available from *http://dx.doi.org/10.6019/EMPIAR-10094*, code from *https://github.com/reyesaldasoro/MitoEM*, and segmented nuclei, cells, invaginations and mitochondria from *https://github.com/reyesaldasoro/HeLa_Cell_Data*.

## 1 INTRODUCTION

The shape of the cellular nuclear envelope is important as it has been linked to processes associated to viral infections [1] as well as cancer [2, 3, 4]. Similarly, mitochondrial function and damage has been found to be correlated to the presence of disease [5, 6], and in cancer in particular the manifold role of mitochondria inside cancerous cells has been previously highlighted [7].

Mitochondria are organelles that can change shape and distribution inside the cell. Moreover, a substantial amount of communication has been shown to exist between mitochondria and the nucleus of a cell [8]. This communication may lead to alterations between the two organelles - nuclear genes may be altered by an impairment in the mitochondria [9]. The damage in both structures can contribute to the progression and metastasis of cancer [10]. A relationship between components of the cytoskeleton and the distribution and shape of mitochondria has been explicitly described multiple times [11, 12]. However, up to the best knowledge of the authors, no explicit relationship has been found between the shape of the nucleus of a cell and the distribution or shape of mitochondria in the same cell.

Electron microscopy (EM) has the capability of creating high-resolution images that can be utilised to analyse the structure of the cell and its organelles. However, computational methods to identify and characterise the structures in order to make inferences are yet being developed to produce high-accuracy results on EM images. This is due to the low contrast and high complexity of the structures revealed by EM imaging. Many techniques have been utilised to perform segmentation on EM images, from traditional image processing [13] to deep learning methodologies [14]. Methods based on Persistent Homology (PH) have shown satisfactory results on other image segmentation tasks [15], but have not been applied as widely as other methodologies to EM images and thus are worth exploring in the context of segmentation of subcellular structures.

In this paper, a methodology to analyse relationships between the nuclear envelope and mitochondria inside HeLa cell images obtained using EM is described. For this goal, four mitochondria segmentation algorithms were compared. These algorithms were based on conventional image segmentation methods (canny edge detection, hole-filling, image morphology), persistent homology, a hybrid algorithm combining the two previous methods, and deep learning. Metrics were extracted from the segmented structures and compared.

## 2 MATERIALS AND METHODS

### 2.1 Materials

Details of the original serial block face scanning electron microscopy (SBF SEM) datasets of *HeLa* cells have been previously described [16] and illustrated in Fig. 1. Nuclear envelope and plasma membranes of 25 cells were segmented from the 8, 192*×* 8, 192*×* 517 voxel dataset [13] and the 3*D* segmentations were used to select the region in outside the nucleus and inside the plasma membrane in subsequent steps of the methodology.

**Fig. 1.**
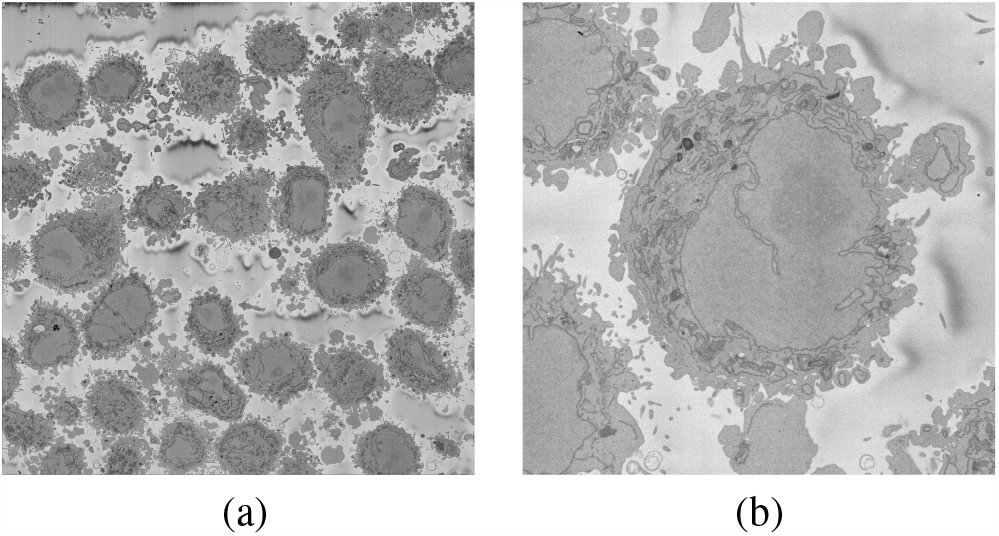
(a) A slice of the original electron microscopy (EM) volume containing HeLa cells. (b) A magnified view of region of interest (ROI) from the original EM slice.

### 2.2 Methods

#### 2.2.1 Segmentation of Invaginations

Invaginations were segmented on a per-slice basis with the following sequence of traditional image processing steps. The nuclear region was filled for holes, morphologically closed with a rather large structural element, and eroded with a rather small structural element to remove the small sections on the surface of the nucleus that did not really penetrate inside the nucleus. To measure how deep the invaginations penetrated into the nucleus, a Euclidean distance map was generated from the region outside of the nuclear envelope, including the filled invaginations. Then, for each invagination, the mean and maximum distance were calculated.

#### 2.2.2 Mitochondria Ground Truth (GT) of Benchmark Set

A benchmark set of five slices belonging to a single cell was used to test algorithms. The mitochondria in the benchmark images were segmented manually (by D.B.-P.) in 2*D* by delineating their perimeter. For comparison, a second segmentation (by C.C.R.-A.) was performed blind to the first segmentation.

#### 2.2.3 Comparison of Mitochondria Segmentation Algorithms

In order to segment mitochondria within individual cells, an evaluation of four distinct algorithms was performed: Persistent Homology (PH), traditional image processing methods, a hybrid algorithm of the previous two, and a deep learning model known as MitoNet [14]. The algorithms were selected based on the average Intersection-Over-Union (IoU) they obtained over the benchmark images.

#### 2.2.4 Topology Segmentation Algorithm

A segmentation algorithm based on Persistent Homology was implemented following the methodology described in [15]. First, histogram equalisation was applied on each of the cytoplasm slices. Upper-thresholds for each value in the range of 0 to 255 were applied to make 256 binary images. Using these thresholded images, the first two Betti numbers *β*_0_ and *β*_1_ were calculated. The maps of threshold value to each Betti number is called the Persistent Homology Profile (PHP). Using these values we can calculate another profile given by the values of *β*_0_*/β*_1_, these are the values that were used to classify each patch as having a part of a mitochondrion or not.

A Random Forest model *M*_*RF*_ was trained on the PHP’s extracted from patches that contained complete mitochondria. The model used 100 estimators and no maximum depth. In order to perform the final classification, the 2*D* slice was split into square patches of varying side lengths (30, 40, 50, 60, 75, 90) with 30% overlap. The PHP was extracted from the patch and *M*_*RF*_ classifies it. From the overlap percentage and the amount of side lengths, it followed that each pixel could be at most part of 24 different patches. Each pixel was then assigned a confidence value between 0 and 24, corresponding to how many patches it belonged to that were classified as a containing part of a mitochondrion by *M*_*RF*_ . Any pixel with a confidence value greater than or equal to 5 was classified as a mitochondria patch.

#### 2.2.5 Traditional Segmentation Algorithm

The segmentation of the mitochondria exploited the fact that the mitochondria have a boundary that is darker than the region that surrounds it and that it is a closed structure. Thus, the region between the nucleus and background was thresholded, then morphologically thinned so that all lines were 1 pixel wide. These regions were filled for holes and then opened with a morphological operator size 3*×* 3. The effect of these steps was that closed regions, like circles or ellipses, would be unaffected, but lines that were not closed would be removed. The threshold was selected to span the dark and bright intensities of the cell and the results of each level were grouped. A confidence parameter was determined by the number of times that regions appeared with this process. Regions that appeared at least twice were selected as mitochondria.

#### 2.2.6 Hybrid Segmentation Algorithm

A hybrid of the PH and the traditional algorithm was implemented to complement their performance. Initially the intersection of the two segmentations was taken. Then, regions predicted exclusively by the traditional algorithm were compared against adjacent regions of the intersection. If the traditionally-segmented region’s area was larger than 25% the area of the intersection region, it was added to the final segmentation.

#### 2.2.7 MitoNet Model (empanada)

The MitoNet neural network model was introduced by R. Conrad and K. Narayan in [14]. It is a model based on Panoptic-DeepLab’s architecture [17] which was pre-trained on CEM1.5m (*https://www.ebi.ac.uk/empiar/EMPIAR-11035/*) and trained on CEM-MitoLab (*https://www.ebi.ac.uk/empiar/EMPIAR-11037/*), these two datasets were curated by the creators of the MitoNet model, which contained a large number of EM images from different types of cells and tissues. The model is accessed through a Python package called *empanada* or as a plugin for napari (*https://napari.org/*) with the same name.

#### 2.2.8 Morphology Metrics

Segmentations obtained with the previous algorithms were saved as individual 2*D* images, assembled as 3*D* structures from which metrics were extracted and figures generated using MATLAB^®^ (Mathworks™, USA). The metrics calculated (all in voxels) for each cell were: volume of the cytoplasm (calculated as the volume contained by the plasma membrane minus the volume of the nucleus), total volume of the invaginations, total number of mitochondria, total volume of the mitochondria, average volume of the mitochondria, and average aspect ratio of the mitochondria. The last metric was defined as the ratio of the major axis and the minor axis of each mitochondrion in 3*D*.

## 3 RESULTS AND DISCUSSION

The segmentation of the invaginations was rather good and although it was evaluated only visually invaginations were clearly distinguished (Fig. 2(a)). The 3*D* structures of the invaginations were rather complicated as can be seen in Fig. 2(b) with some superficial invaginations and others that penetrated deep inside the nucleus. It should be noted that the metrics only capture partly the complexity of the invaginations of the nuclear envelope and warrant further study.

**Fig. 2.**
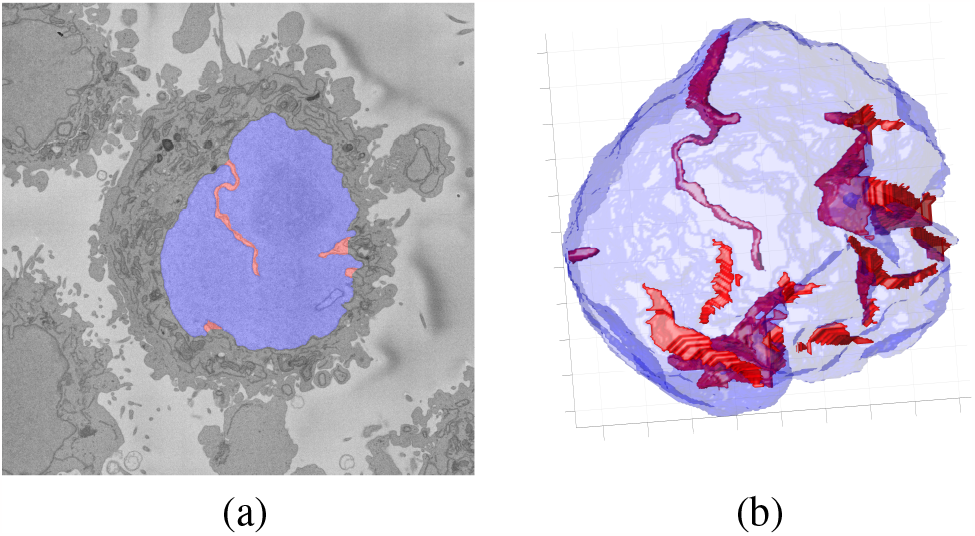
(a) 2*D* segmentation of cell nucleus (blue) and invaginations (red). (b) 3*D* reconstruction of the cell nucleus (blue) and invaginations (red).

The results of the four algorithms are illustrated in Fig. 3 and the numeric results shown in Table 1 and range between 0.41 and 0.65, which might be considered low, but the inter-observer similarity is not much higher at 0.696. This indicates that two experienced observers, who can scroll up and down slices before reaching a decision, do not agree completely and thus the ground truth may not be perfect. MitoNet outperformed the other algorithms by a fair margin and thus selected for subsequent steps. However, it must be noted that the combination of the PH and image processing improved results of the individuals, thus a further development of a single algorithm with elements of both could provide better results.

**Table 1.**
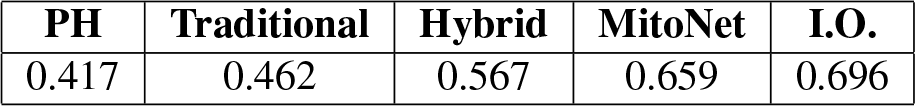
Segmentation results of the methodologies: Persistent Homology (PH) Segmentation, Traditional Segmentation, Hybrid Segmentation MitoNet Model, and Inter-Observer Segmentation (I.O.). Values correspond to Intersection over Union or Jaccard Similarity Index.

**Fig. 3.**
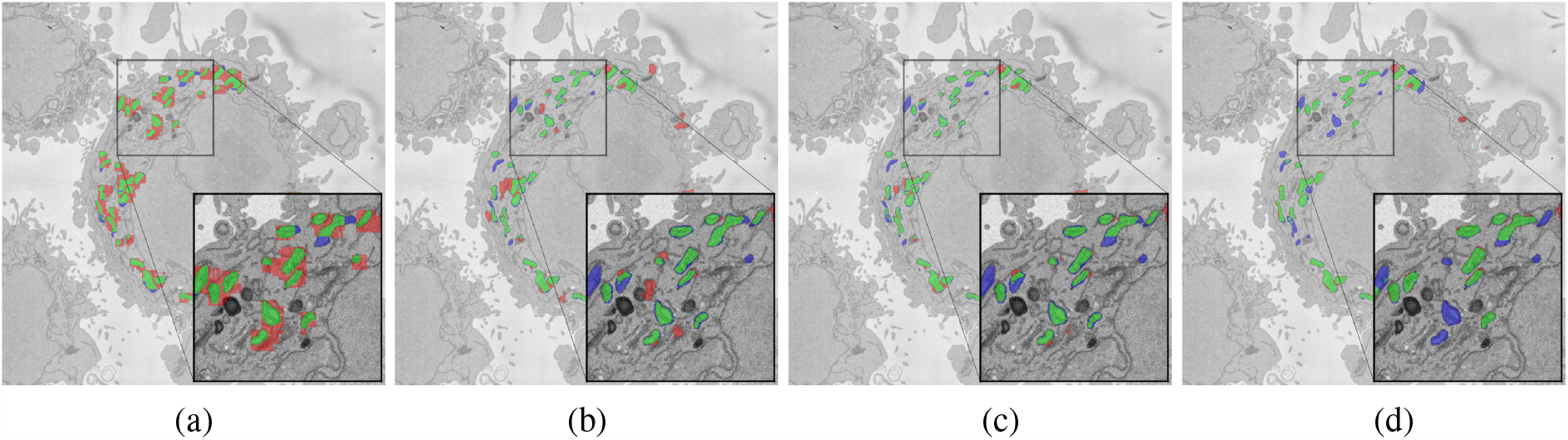
2*D* segmentation of mitochondria on a single cell slice obtained using (a) Persistent Homology (PH) Algorithm, (b) Traditional Segmentation Algorithm, (c) Hybrid Segmentation Algorithm, (d) MitoNet Model. Green pixels show True Positive (TP), red pixels show False Positive (FP), and blue pixels show False Negative (FN). Region of interest (ROI)’s have been darkened to improve contrast.

The 3*D* segmentation of the nucleus, plasma membrane, invaginations and mitochondria is illustrated in Fig. 4. Four cells were selected to represent the variability found within the cells. There were cells where mitochondria were distributed around the nucleus (Fig. 4(a)), concentrated in two extremes suggesting a polarity (Fig. 4(b)), distributed along the nucleus except for a small region (Fig. 4(c)) and concentrated on just one side of the cell (Fig. 4(d)). The variability of these results are consistent with the literature [11]. A natural extension of these observations is to see the cells in the presence of their neighbours and not in isolation.

**Fig. 4.**
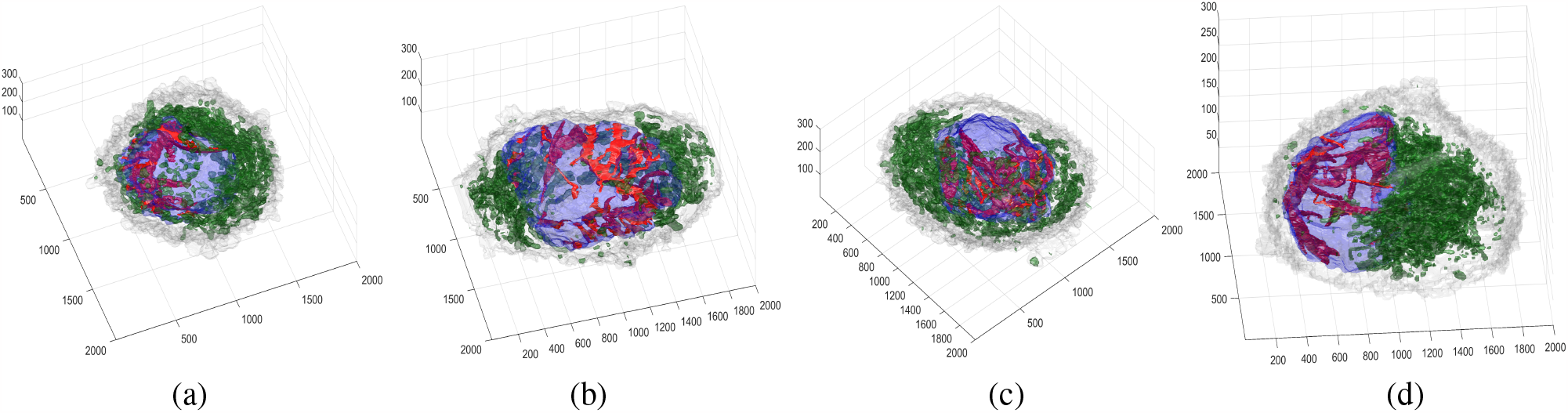
Illustration of the variability of the distribution of the cellular structures in four different cells. In all cells, the plasma membrane is displayed in a very light gray, the nuclear envelope in light blue, invaginations of the nucleus in bright red, and mitochondria in green. It can be noticed how mitochondria distribute in the cells: (a) uniform, (b) polarised towards left and right, (c) uniform except for a small region, (c) concentrated towards the right.

Pearson correlation coefficients were calculated pair-wise between the morphological metrics previously described and the following results were interesting:

**1** A positive (*r* = 0.5067) and significant (*p* = 0.0097) correlation between the total volume of invaginations and the total volume of mitochondria. Whilst this could suggest that larger and more complex invaginations were associated with more mitochondria, the positive correlation may just be an indication of the size of the cell.

**2** A negative (*r* =*−* 0.4466) and significant (*p* = 0.0252) correlation between the number of mitochondria and the average volume of mitochondria was found. This suggests that the more mitochondria are present in a cell, the smaller they are. This correlation would not be affected by the size of the cell.

**3** A negative (*r* = *−*0.4407) and significant (*p* = 0.0275) correlation between the volume of the cytoplasm and the aspect ratio of the mitochondria, suggesting that the larger the cytoplasm, the thinner and more elongated the mitochondria.

## 4 CONCLUSION

In this paper, a methodology to morphologically analyse HeLa cells as observed with Electron Microscopy has been described.

In order to characterise the shape of the nucleus, the invaginations of the nuclear envelope were segmented using conventional image processing methods. A segmentation of mitochondria was also performed, during this process, different algorithms were compared in order to acquire a segmentation that yields results similar to manual segmentation. The deep learning model known as MitoNet outperformed all other algorithms and was chosen to perform such segmentation. PH, hybrid and MitoNet were implemented in Python. Image processing, the invaginations and metrics and correlations were measured using MATLAB®.

The measurements were compared and correlations were found between the total volume of invaginations and the total volume of mitochondria (*r* = 0.5067), the total volume of the cytoplasm and the aspect ratio of the mitochondria (*r* =*−* 0.4407), finally, the number of mitochondria and the average volume of mitochondria (*r* =*−* 0.4466). Whilst these results are interesting, it is acknowledged that much more could be explored in the future. Specifically, improving the segmentation of mitochondria, extracting more morphological measurements and developing the study of the correlations between metrics.

## Acknowledgements

The authors would like to thank Dr Martin L. Jones and Dr Lucy M. Collinson from The Francis Crick Institute for their valuable input in this work.

